# Hyperactive PI3Kinase delta enables long distance regeneration of the rat corticospinal tract

**DOI:** 10.1101/2023.10.27.564182

**Authors:** Kristyna Karova, Zuzana Polcanova, Stepanka Suchankova, Lydia Knight, Bart Nieuwenhuis, Radovan Holota, Vit Herynek, Lucia Machova Urdzikova, Rostislav Turecek, Jessica C.F. Kwok, Joost Verhaagen, Richard Eva, James W Fawcett, Pavla Jendelova

## Abstract

Maturation of central nervous system neurons leads to loss of their intrinsic regeneration potential. In particular after injury of the adult spinal cord there is minimal regeneration of corticospinal axons, which control gait and fine movement. Previous work has shown that knockdown of PTEN to increase PIP3 levels can promote regeneration in young animals, but the effect is much less in adults probably due to low PIP3 production. Here, we have transduced sensorimotor cortex neurons with a hyperactive form of PI3K, PI3Kδ, which increases PIP3 in mature neurons. This enables cortical neurons to regenerate corticospinal axons and improve behavioural outcomes.

We used a C4 dorsal column lesion model in adult rats and injected the right motor cortex at 4 sites concurrently with a mixture AAV1-PIK3CD and AAV1-eGFP or titre matched AAV1-eGFP only. We allowed rats to survive for 6, 9, 12 or 16 weeks. Immunostaining showed 70 - 80% co-expression in cortical neurons which remained stable at both 12 and 16 weeks. We counted GFP labelled axons in 20 μm spinal cord sections. In PI3KCD-treated animals many axons were seen to have regenerated around the margins of lesions, collecting into a knot of axons with the typical appearance of regeneration at the caudal end. Tracing down the cord, and excluding axons and neurites that could have come from unlesioned ventral CST, we found axons extending up to 1 cm below lesions, numbers decreasing with distance from the lesion. After 16 weeks there were circa 200 axons at the caudal end of lesions with a regeneration index of 0.2, with half this number at 12 weeks. Behavioural testing for 16 weeks revealed functional improvements in skilled paw reaching, grip strength and ladder rung walking in rats treated with PIK3CD compared to GFP only controls. In addition to behavioural testing, functional recovery of PIK3CD treated rats was confirmed with electrophysiological recordings during which we stimulated the right pyramid. Cord dorsum potentials (CDPs) above and below lesion and EMG forepaw distal flexor muscles showed greatly increased connectivity compared with GFP only controls, lesion only controls and uninjured shams. We conclude that forcing upregulation of PI3Kδ in cortical neurons leads to robust regeneration after spinal cord injury that results in functional restoration.

## Introduction

The intrinsic ability of axons in the brain and spinal cord to regenerate diminishes as neurons mature. There are several mechanisms behind this loss of regenerative ability, notably 1) reduction in signalling particularly in the PI3K/AKT pathway, 2) development of neuronal polarity, leading to polarized transport that excludes many growth-related molecules from axons, 3) genetic/epigenetic changes in maturing neurons that affect expression of axon growth-related molecules.

A key signalling pathway for the control of cell movement and migration is PI3kinase/PIP3. Unlike PIP2 which is widespread, PIP3 levels are tightly controlled and localized through control of the activity of PI3K and degradation of PIP3 via PTEN and SHIP [1]. PIP3 produced by PI3K stimulates cell motility, protein translation, transport and other functions via signalling through or in concert with the Akt/mTOR, /GSK3, CRMP2, NFκB pathways [2], transport via Arf GAPs and GEFs, and through various molecules with PH and FYVE domains [3]. It has been shown, that axon growth is associated with an increase of axonal PIP3 both during development and regeneration [4], and manipulation of PI3K to mediate PIP3 increase prevented growth cone collapse in DRG sensory neurons while PTEN facilitated it [5]. The involvement of PIP3 in axon regeneration has been confirmed in many experiments that have demonstrated enhanced axon regeneration in young rodents after PTEN deletion [6][7][8]. Other studies have revealed, that stimulating the PI3K/Akt/mTOR axis results in axon regeneration after spinal cord injury. In one such study, IGF-1 used together with osteopontin achieved robust short distance regeneration having activated the PI3K/Akt/mTOR pathway and the integrin domain [9].

PIP3 levels are high in immature axons but drop to a low level as they reach their targets and mature [4]. PIP3 can be increased in young neurons by reducing its degradation through deletion of PTEN, but this becomes less effective with greater maturity. Neuronal levels of PI3Kα are high, so it is probable that activation of PI3Kα is reduced with maturity, possibly because many PI3K-activating receptors are excluded from mature axons. This makes PTEN knockdown less effective in mature neurons [10]. In order to increase PIP3 levels in mature neurons, a hyperactive form of PI3K is needed. PI3Kδ is a hyperactive form normally found in the immune system and sensory neurons [11]. Expression of this enzyme in cortical neurons enabled vigorous regeneration in an *in vitro* axotomy assay [4], with a dose-related effect, and expression in retinal ganglion cells enabled regeneration of their axons in the adult mouse optic nerve. Transport of integrins and recycling endosomes in which they travel was also restored into mature axons, probably due to the effects of PIP3 on GEFs and GAPs that affect endosomal transport.

In the present study we have asked whether expression of PI3Kδ in neurons of the rat sensorimotor cortex can enable regeneration of axons of the corticospinal tract. We show that many axons are able to regenerate for distances of up to 1 cm, with restoration of sensorimotor behaviour and physiological connections to forelimb muscles.

## Methods

### Animals

All experiments were performed in accordance with the European Communities council directive of 22^nd^ of September 2010 (2010/63/EU) and were approved by the Ethics Committee of the Institute of Experimental Medicine ASCR, Prague, Czech Republic.

Young adult male and female Wistar or Lister Hooded rats were used in this study. Rats were housed in pairs with a 12h light/dark cycle and provided with water and food *ad libitum*. They were quarantined for 2 weeks prior to surgery after which they were checked every day with increased care taken for the first 3-5 days and food pellets placed inside their cage.

### Viral vectors preparation

Recombinant AAV vectors were produced as previously described [12]. cDNA constructs were designed and obtained from Addgene, scaled in house using DH5α transformation competent cells (Invitrogen, Waltham, MA, USA) and isolated with Maxiprep (ThermoFisher, Waltham, MA, USA). Subsequently, correct ITR sites were verified with a restriction analysis and electrophoresis. Plasmids were then used to generate AAV1-CAG-PIK3CD, titre 5 × 10^12^ gc/ml; AAV1-CAG-eGFP, titre 2.64 × 10^12^ gc/ml; AAV1-hSYN-PIK3CD, titre 2.7 × 10^12^ gc/ml; AAV1-hSYN-eGFP was purchased from Vigene with a titre of 2.3 × 10^13^ (now distributed by Charles River, cat. number CV17001-AV1). A mixture of AAV1-CAG-eGFP : AAV1-CAG-PIK3CD, AAV1-hSYN-eGFP : AAV1-hSYN-PIK3CD at a ratio 1:10 or titre matched AVV1-hSYN-eGFP alone were used.

### Spinal cord injury and virus injections

Rats weighing 220 – 400 g were anaesthetized with 3% isoflurane, shaved and placed on a heating pad that was kept at 37°C, their temperature monitored with rectal thermometer. At this stage, buprenorphine (Bupaq Multidose 0.3 mg/ml, Richter Pharma, Austria, 0.01 mg/kg, s.c.) and caprofen (Rymadil, Pfizer, 7.5 mg/kg, i.m.) were administered as pain relief. An incision was made in the neck area and through cervical muscles to gradually expose the spine with blunt muscle retraction used whenever possible. Laminectomy of the C4 vertebra was performed and small punctures into the dura mater were made with a needle. A set of micro forceps (0.25 mm tip, Fine Science Tools, Foster City, CA, USA, 11083-07) was inserted into the punctures to a depth of 1 mm and carefully but firmly squeezed for 7 seconds. Animals were then sutured in anatomical layers and skin was treated with Novikov solution.

Within the same surgery, 4 injections with either injection water (aqua ad inj.), AAV1-hSYN-eGFP, AAV1-hSYN-eGFP + AAV1-hSYN-PIK3CD or AAV1-CAG-eGFP + AAV1-CAG-PIK3CD. Animals were fixed onto a semiautonomous stereotaxic frame (Neurostar, Tubingen, Germany). The scull was exposed, positions of drill and syringe were synchronized and calibrated relative to bregma and lambda. Corrections for tilt and scaling were made before drilling. Each site was then injected with 0.5 μl mixture at rate of 0.2 μl/min at 1.5 mm depth using a 10μl Hamilton syringe. Coordinates used were as follows: ML/AP: 1’5, 1; 2, 3’5; 3’5, 2; 3, 0’5. Needle was left in place for 3 minutes after injection and retracted in 0.5 mm increments every 30 s. Skin on the scalp was then sutured and treated with Novikov solution. Each rat received 2 ml of glucose solution (Ardeanutrisol, Glucose 100g/l, Ardeapharma, Sevetin, Czech Republic) at the end of surgery.

### In vitro MR imaging of the spinal cord sections

Spinal cord sections were measured immersed in PBS solution in 2 ml test tubes. MR images were obtained using a 7T MR scanner MRS*DRYMAG 7.0T (MR Solutions, Guildford, UK) equipped with a mouse head resonator coil. Three sequences with high resolution were acquired. A T2-weighted turbo-spin echo sequence in axial direction, repetition time T = 4000 ms, turbofactor TF = 8, echo spacing TE = 8 ms, effective TE = 40 ms. The number of acquisitions AC = 16 with acquisition time approx. 17 minutes. Acquired matrix was 128x, field of view FOV = 10 × 10 mm2, 20 slices with slice thickness 0.5 mm with no gap. A T1-weighted axial images were obtained using a 3D gradient echo sequence, TR = 10 ms, flip angle 20°, TE = 3.3 ms, number of AC = 16, acquisition time 11 minutes. Matrix was 128 × 128 × 32, FOV = 10 × 10 × 16 mm3 (which provides axial slices with thickness of 0.5 mm).

### Behavioral testing

Lister Hooded rats treated with AAV1-hSYN-GFP + AAV1-hSYN-PIK3CD (n=15) or AAV1-hSYN-GFP (n=14) only were tested with a range of behavioral tests to assess changes in functional motor and/or sensory function with a focus on front limbs. Rats were trained for two weeks daily before surgery to be able to perform skilled paw reaching, grip test, ladder rung walking. After this period, baseline measurements were recorded for all tests. After surgery and vector injections, rats were allowed 1 week recovery break and then tested once a week for 16 following weeks.

#### Skilled paw reaching

To determine improvements in fine motor control, animals were placed into a Montoya staircase and allowed to retrieve sugar pellets for 15 minutes. At the start of the test, each stairwell contained 3 sugar pellets. After 15 minutes, remaining pellets were counted and the longest distance from which a pellet was retrieved was recorded.

#### Grip test

Grip test was used to evaluate muscular strength of rats. They were encouraged to hold on to a grid attached to a force measuring device (BIO - GS3, Bioseb) with both forepaws. They were then pulled back in a horizontal plane until they lost their grip. The peak force applied to the grid was recorded just before the loss of grip. This was repeated 5 times and values were noted. Only 3 highest values were averaged and used in analysis.

#### Ladder rung walking

A ladder with unevenly spaced rungs was used and rats were recorded traversing the ladder 5 times. Animals were encouraged to cross with sugar pellets and their stepping was evaluated according to [13]. Each score was then normalized to the number of steps evaluated per cross over.

#### Von Frey sensory test

The Von Frey test was used to determine differences in sensitivity to mechanical noxious stimuli. The device used consisted of transparent compartments where rats were placed standing on a mesh platform through which Von Frey fibers were applied to the plantar surface of each forepaw (IITC IncLife Science, CA, USA). Prior to the test, rats were placed into the compartments and allowed to acclimatize for 15 minutes. Each forepaw was measured 5 times and the value upon paw withdrawal was recorded. Care was taken to ensure the rat was unaware of the fiber being applied. For statistical analysis, the upper and lower extreme values were first omitted before averaging the remaining 3 values.

### Electrophysiology recording

In a terminal setting, Wistar rats treated with PI3Kδ (n=11) or GFP vectors (n=7) surviving for 16 weeks after injury and treatment, and uninjured controls (n=5) were anaesthetized with urethan (1.5 g kg^-1^). An incision was made in the neck and back areas to gradually work through the muscle layers and expose the obex by removing the cerebellar tonsil as described previously [14,15]. Subsequently, the cervical spinal cord including the lesion and up to 1 cm below was also exposed. A stimulating tungsten electrode with impedance 100 kΩ was lowered into the right pyramid. Concurrently, a stationary silver ball electrode was placed on the cord surface 1 cm cranially to the lesion site as a control of correct stimulating electrode placement. Another silver ball electrode was placed 1 cm below lesion 1 mm laterally from midline to the left. Silver ball electrodes were used to measure cord dorsum potentials (CDPs). Within the same setting, subcutaneous silver electrodes were inserted into the distal forearm muscles of both left and right forearms for electromyography (EMG) recordings. Potentials were evoked via stimulation with five square-wave pulses at 300 Hz, 30-300 μA, 400 μs, delivered every 0.33 s as used previously [16]. Responses were recorded 3-5 times with each response approximating 50 stimulations. Once all measurements were completed, a re-lesion was performed to confirm loss of response and to validate recordings. Rats were then maintained for 2.5 hours to allow for cFOS upregulation.

### Imunohistochemistry

Floating 40 μm frozen brain coronal sections or 20 μm mounted spinal cord cross or 20 μm sagittal/frontal sections were washed with TRIS buffered saline (TBS). When staining for PI3Kδ, heat induced antigen retrieval (HIER) step was incorporated into the protocol. Sections were washed once in dH2O and transferred into a 4.5 mM aqueous solution of citraconic anhydride. Plates containing the samples were placed in a water bath and warmed to 98°C for 20 mins. Samples were then left to cool completely before washing once in dH_2_O and twice in TBS before continuing with standard protocols. Work with the HIER agent was performed in a fume hood. Sections were then permeabilized with TritonX-100 for 20 mins and endogenous avidin/biotin were blocked with Avidin/Biotin block (Abcam, Bristol, UK, ab64212) in samples where biotin-streptavidin amplification was used before blocking with 10% chemiblocker (Merck Millipore, Billerica, MA, USA, #2170) for 2 hours at room temperature. Following incubation in primary antibody, samples were washed and incubated in their respective secondary antibody (1:400, Life Technologies, Carlsbad, CA, USA) or biotinylated secondary antibody (1:400, Vector Biotechnologies) for 2 hours at 4°C, then washed and incubated in Streptavidin Alexa Fluor 488/594 (1:400, Life Technologies, Carlsbad, CA, USA) at 4°C for additional 2 hours and DAPI (1:3000) for 10 mins. After washing, sections were mounted with Vectashield fluorsafe (Vector Laboratories, CA, USA). Primary antibodies used in this study were rabbit anti-p110 (1:300, Abcam, Bristol, UK, ab1678), chicken anti-GFP IgY fraction (1:400, Thermo Fisher Scientific, Waltham, MA, USA, A10262), mouse anti-pS6 (1:300, Cell Signaling, Danvers, MA, USA, #62016), mouse anti-PKCγ (1:1000, Santa Cruz, Dallas, TX, USA, sc166451 A-7 clone), mouse anti-cFOS (1:500, Abcam, Bristol, UK, ab208942), rabbit vGlut1/2 (1:500, Synaptic Systems, Göttingen, Germany, #135503). Images of cross sections 1 cm below lesion to visualize vGlut1/2 together with GFP were acquired with Andor Dragonfly 503 spinning disc confocal microscope (Oxford Instruments, Abingdon, UK).

#### Co-expression analysis and soma size

Co-expression of PI3Kδ and GFP was determined in 4-5 brain coronal sections per rat 12 (n=4) and 16 (n=8) weeks after surgery using the Imaris 9.4 software. Images were obtained using a confocal microscope at 20x magnification (Olympus, SpinSR10). First, PI3Kδ^+^ cells were identified from which a PI3Kδ^+^GFP^+^ fraction was determined.

Using ImageJ (NIH, Bethesda, MD, USA), neural soma length was determined in 5 sections per rat in PI3Kδ expressing neurons after 12 weeks (n=4) and 16 weeks (n=4) and compared with lengths from respective control groups expressing GFP only (n=3 and 3) One-way ANOVA with Tukey post hoc tests were used.

#### Axon sprouting analysis

To quantify the number of axons sprouting above the lesion following C4 dorsal column crush, we used spinning disc (20x magnification) to image cervical spinal cord cross sections (5-6 sections per animal) at 100 μm increments. The number of GFP+ axons were counted using ImageJ, by an observer blind to experimental groups. Briefly, a horizontal line was drawn from the central canal to the lateral grey and white matter border, then an average of 10 vertical lines spaced 100 μm apart were used to count axons in the contralateral grey matter of each 100 μm section. Differences in the number of axons in ventral and dorsal horns was also observed, and all values normalised to the MFI of the dCST (average from 3 images per rat) and area of dorsal/ventral horns. We also analysed cross sections from rats injected with vectors carrying transgenes under the CAG promoter. The number of GFP+ axons was averaged across the 5-6 images and compared between experimental groups (n= 3-4 per group) using GraphPad prism software, presented as mean ± SEM. One-way ANOVA with Tukey post hoc tests were used.

#### Axon counting and regeneration index

Every other sagittal 20 μm section was used for staining against GFP and axon counting in PI3Kδ treated rats and their controls 12 weeks (n=4 and 3) and 16 weeks (n=4 and 5) after spinal cord injury and cortical injections. A grid with 60 μm spacing placed in microscope eyepiece allowed to determine the distance bellow lesion. A count of axons every 60 μm was determined using Axioskop 2 plus microscope (Zeiss, Oberkochen, Germany). At each 60 μm increment, the number of axons crossing a line was noted. Axons appearing to be part of the ventral CST, which are not destroyed in this model were not counted. Axons clearly sprouting from the ventral CST were also not included. For plotting reasons, sums of axons of every 600 μm increment were used, which were a product of a sum of all sections counted that corresponded to the same caudal distance from lesion. A regeneration index was then calculated as the number of counted axons normalized to the mean number of axons in the dorsal CST counted in 3 cross sections above the lesion. Paired t test was used to calculate significant differences between groups. A reconstruction of regenerated spinal cord was produced by approximate drawing of GFP^+^ axons and neurites. Blue color was assigned to those found cranially to lesion and extending, red colour was assigned to axons and neurites in the ventral part of sections and green colour represents neurites residing in the dorsal half of the cord caudal to the lesion border (Fig. 4C).

#### cFOS analysis

Spinal cross sections from rats examined in the electrophysiology part of this study (PI3Kδ treated n=11, GFP controls n=5, uninjured n=5) from 0.5 cm above lesion site and 1 cm below lesion were stained for cFOS, an early marker of neuronal activity [17]. Three rats were excluded from analysis, because they had to be euthanized before they were suitable for cFOS analysis due to the required survival time after stimulation. Images of 3 sections per rat were taken with an epifluorescent microscope Leica CTR6500 equipped with a TissueFAXS 4.2.6245.1020 software (TissueGnostics, Vienna, Austria) and were then analysed with ImageJ in 5 regions – central canal area (CC), contralateral ventral horn (VH1 contra), contralateral dorsal horn (DH1 contra), ipsilateral ventral horn (VH2 ipsi), ipsilateral dorsal horn (DH2 ipsi). The number of cFOS^+^ nuclei was then normalized to the selected area size. One-way ANOVA or unpaired t tests were used in statistical analysis.

## Results

### PI3Kδ co-expressed with GFP at high levels and prevented neural soma size reduction

First, we assessed PI3Kδ and GFP protein co-expression in layer V cortical neurons. AAV1 injections led to expression of PI3Kδ and GFP in injection sites with a soma length of around 20 μm. First, the number of PI3Kδ^+^ neurons was determined, from which a percentage of coexpressing neurons emerged by establishing how many of them were also GFP^+^. With the AAV1 mixture that we used, we observed on average 79.7% co-transduction at 12 weeks post-surgery and 68.92% at 16 weeks post-surgery, which indicated that this method was suitable for axon tracing (Fig.1A, B, C). It has been reported, that stimulating PI3K/pAkt/mTOR pathway with high titer vectors can lead to disproportionate soma size increases that were associated with several negative effects [18]. We therefore measured soma lengths of neurons in injured PI3Kδ treated rats and GFP controls. We found that spinal cord injury led to reduction in soma size in GFP expressing controls neurons, which was mitigated by the PI3Kδ treatment. Neurons expressing PI3Kδ fell into a normal soma size range of young adult rats ([19] (Fig.1D).

### PI3Kδ increased pS6 expression in transduced cortex

After transduction with AAV1-hSYN-PIK3CD, we observed increased immunostaining for phosphorylated S6 protein within injection sites, which is a downstream effector of the PI3K/pAkt/mTOR pathway (Fig. 2A, B). This increase persisted through both 12 and 16 weeks with higher levels at 12 weeks and a slight decrease at 16 weeks (Fig. 2C).

**Fig. 1.**
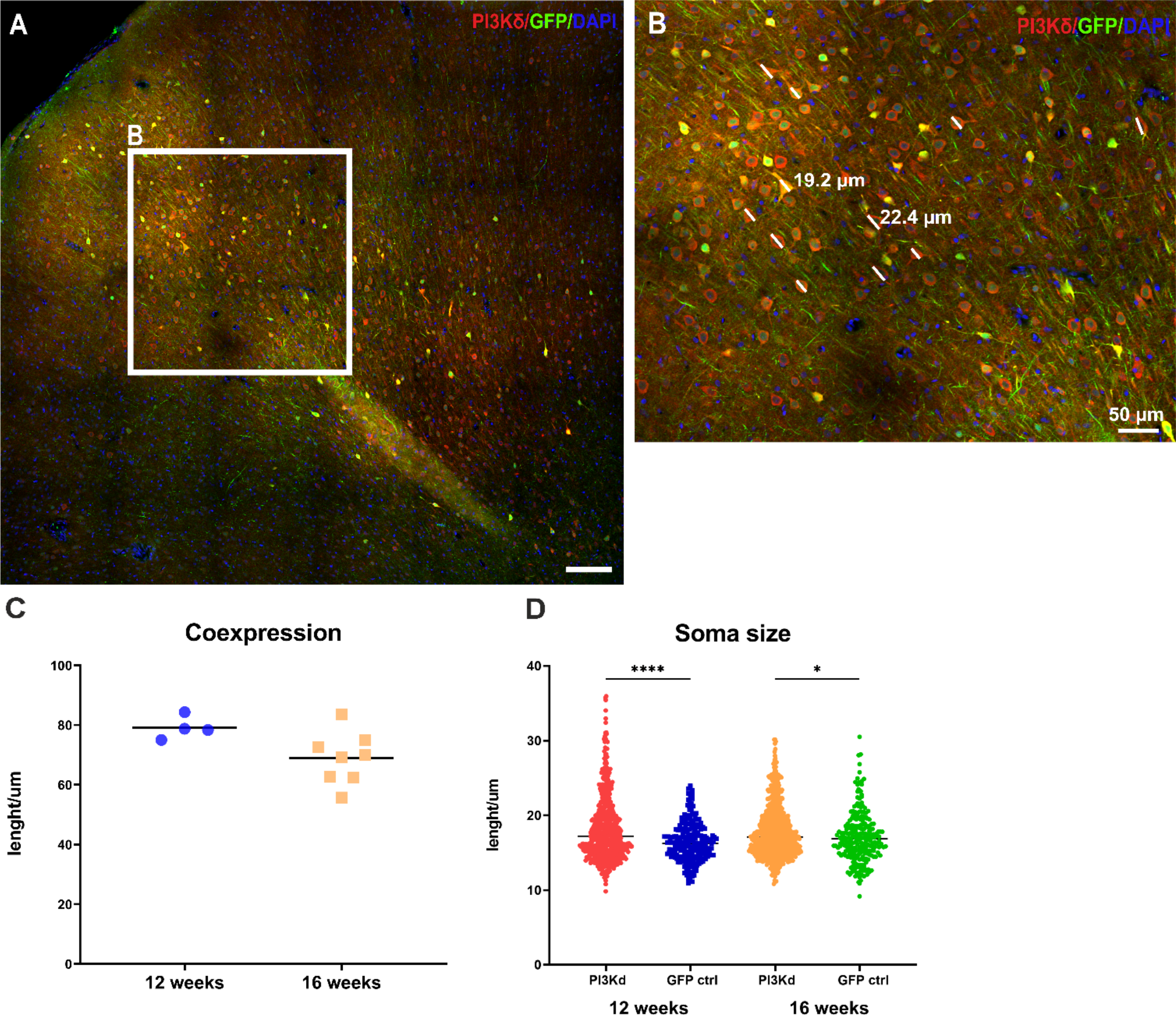
Transduced brains were stained against PI3Kδ and GFP 12 weeks (A, B) and 16 weeks after surgery to determine co-expression of PI3Kδ^+^ layer V neurons positive for GFP tracer. On average, co-expression levels were close to 80% at 12 weeks and declined slightly to around 70% at 16 weeks after cortical injections (C). Lengths of layer V cortical neurons were measured (B, D). At both 12 and 16 weeks after SCI, soma lengths of neurons in control groups were smaller than those overexpressing PI3Kδ (D). Magnification 40x, scale bar 200 μm unless indicated otherwise. One-way ANOVA, **** *p<0*.*0001*, * *p<0*.*05*

**Fig. 2.**
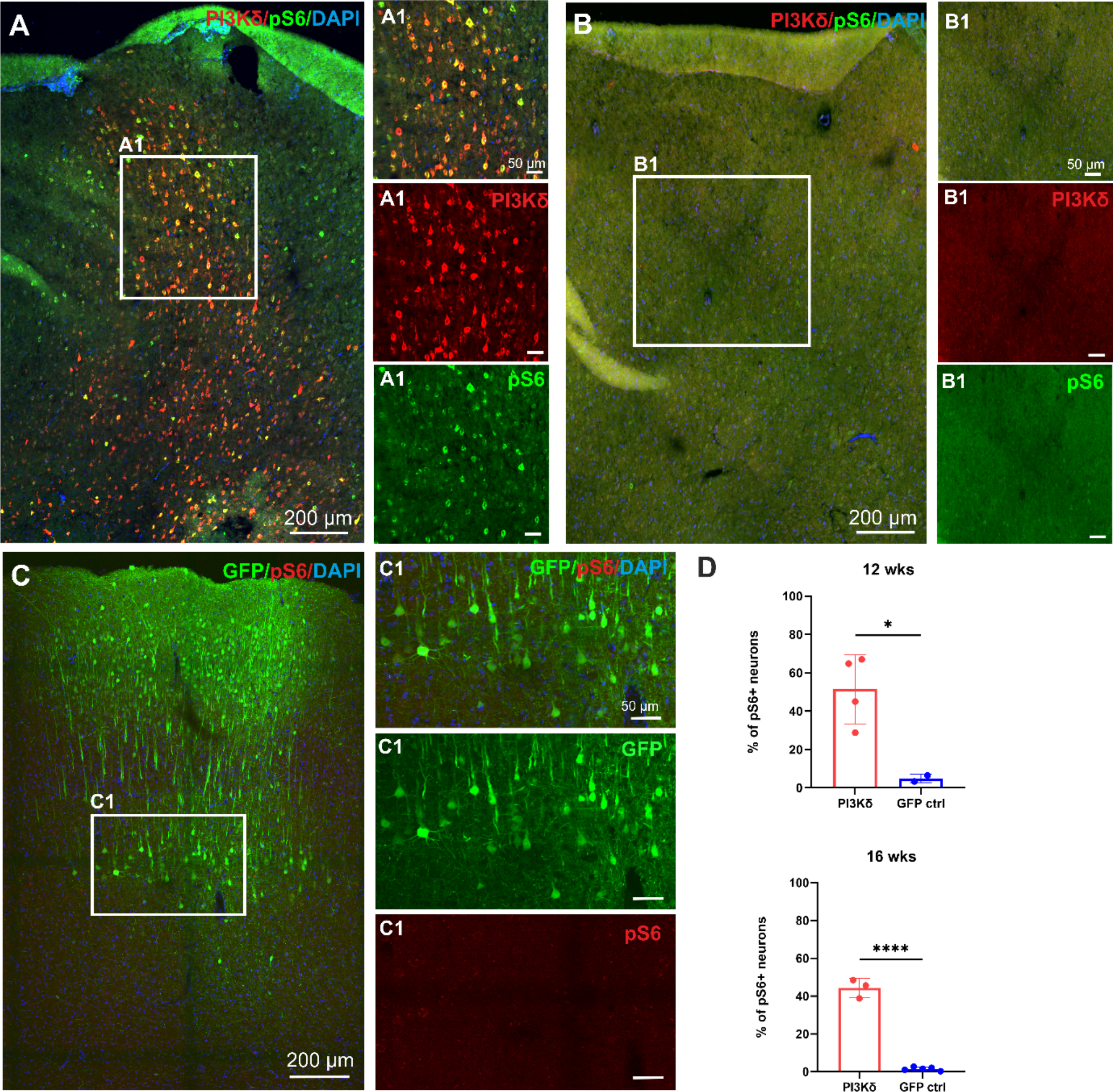
Phosphorylation of S6 remains upregulated in PI3Kδ overexpressing neurons 16 weeks after treatment (A, A1) suggesting active PI3K/pAkt/mTOR pathway. Very low pS6 levels were detected in noninjected cortices (B, B1) and cortices injected with control AAV1 (C, C1). Quantification of of pS6 coexpression with PI3Kδ or GFP shows significant increase of S6 phosphorylation at both 12 and 16 weeks after cortical injections (D). Magnification 20x. Data shown as mean and SEM, Student’s t test, *\* p=0*.*0266, **** p<0*.*0001*

### PI3Kδ treatment increased sprouting of axons across midline above the C4 lesion

Next, we wanted to determine whether PIK3CD treatment induces sprouting of axons in areas above the lesion, where they are intact. We analysed cross sections and counted axons that have sprouted across the midline, and were present contralaterally to the labeled dorsal corticospinal tract (dCST) (Fig. 3A) 12 weeks and 16 weeks after treatment (Fig. 3B, B1) and compared that to GFP only control at the same time points (Fig. 3C, C1). There was extensive sprouting of axons across the midline in the treatment groups, with many more at 16 weeks (Fig. 3Da). Small numbers of shorter neurites were found in the control groups (Fig. 3C, C1). We also analysed the distribution of axons and found that the majority of axons entered the intermediate and ventral horn area. (Fig. 3Db). Additionally, we compared sprouting power of treatment vector with CAG promoter and found more neurites at 12 weeks than we did at 9 weeks (Fig. 5Ba). Additional comparison between either neuron specific hSYN and CAG promoter showed similar numbers of axons crossing the midline in either group at 12 weeks after treatment (Fig. 5Bb). This shows, that PI3Kδ was effective under either of the promoters.

**Fig. 3.**
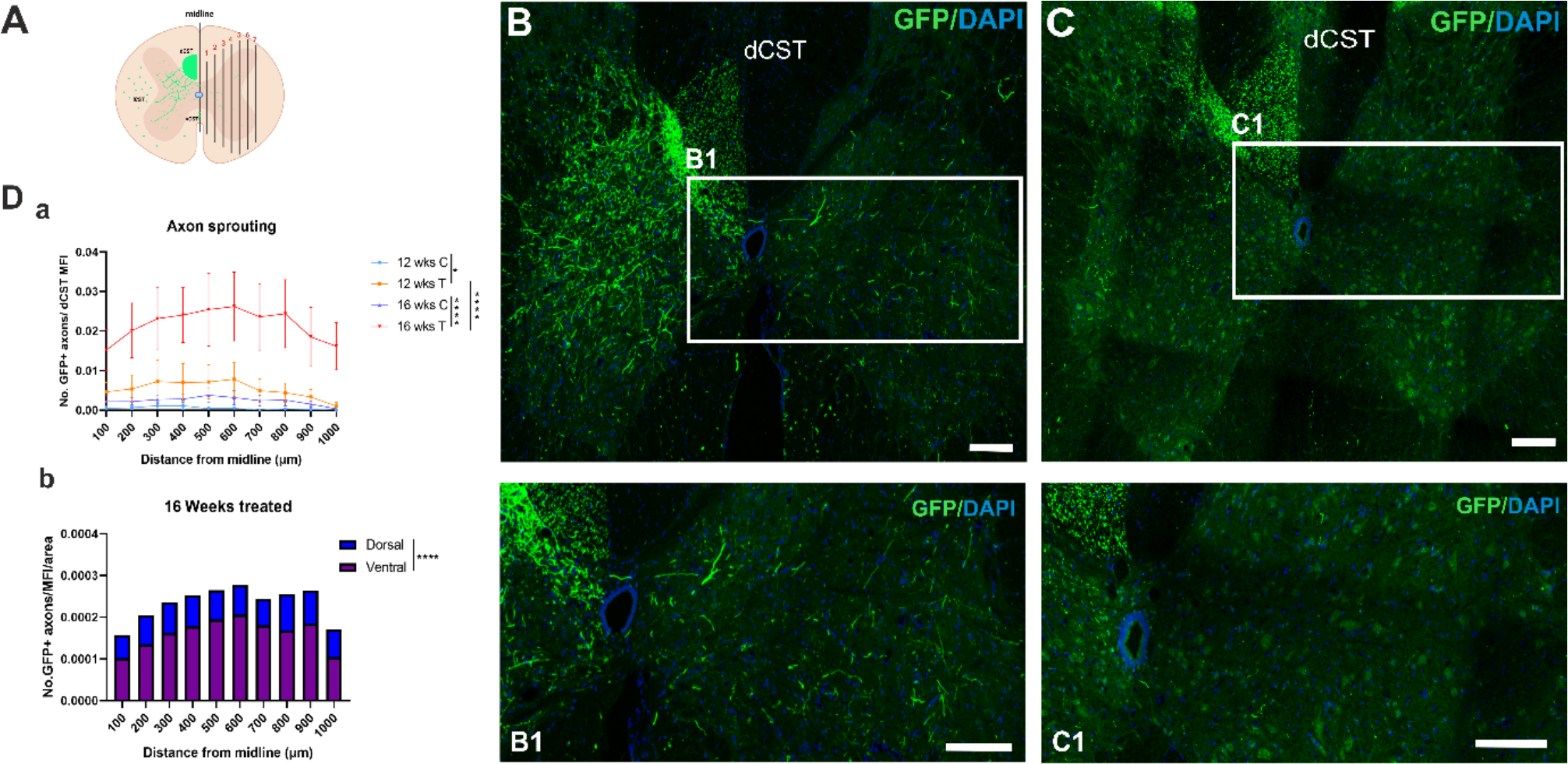
Axon sprouting was determined by counting the number of axons in cervical spinal cord above lesion crossing the midline at 100 μm intervals (A, Da). Distribution of axons in dorsal and ventral horns was calculated (Db) Representative images of spinal cords from treated animals 16 weeks (B) after SCI and control at 16 weeks after SCI (C). Scale bars 200 μm. Data shown as mean and SEM, One-way ANOVA, *\* p=0*.*019, *** p<0*.*001, **** p<0*.*0001*, dCST dorsal corticospinal tract.

### Overexpressing PI3Kδ lead to the growth and sprouting of many axons over long distances below the C4 lesion

To evaluate the number of axons that grew below the lesion site, we counted them in every other 20 μm sagittal section. To ensure that only complete lesions were analyzed, we performed MRI imaging on whole spinal cord segments and PKCγ staining on cross sections from lumbar spinal cords (S. 1). Lesions produced a cavity in the dorsal cord reaching down to or beyond the central canal, roofed by a fibroblastic/meningeal structure. At the rostral end of lesions there were many transected axons, which were in contact with the lesion edge. In treated animals many cut axons could be seen to have sprouted processes that grew around the lesion margins ventrally and laterally. The peri-lesional regeneration was progressive, with few axons at 6 weeks, and progressively more at 9, 12 and 16 weeks (Fig. 4, Fig. 5A, C).

In GFP controls few axons sprouted (Fig. 4B, B1, B2). A reconstructed lesion from a AAV1-hSYN-PIK3CD treated rat is shown in Figure 4C. At the caudal margin of lesion, a knot of regenerated axons was seen as the perilesional axons collected (Fig. 4A, C). From this knot, many axons are seen to grow on in the dorsal cord (Fig. 4A1, A3). These had the typical thin wiggly morphology of regenerates. A progressively decreasing number of these axons was seen to 1 cm below the lesions. The number of axons at the caudal lesion margin was approx. 200 at 16 weeks, 100 at 12 weeks. Comparing labelled axon numbers rostral and caudal to the lesions the regeneration indices were 0.2 and 0.1 respectively (Fig. 4D). A complication in quantifying regeneration in dorsal column lesions is that the ventral CST is not cut, and the axons of this pathway may sprout after injury. It is easy to distinguish the unlesioned axons because they are straight with a large diameter compared to regenerates (annotated with red arrows in Fig. 4A). However, the sprouts are difficult to distinguish from regenerated axons. This is illustrated in Figure 4C, where sprouted ventral axons are shown in red. The position of regenerated CST axons in the dorsal cord, their continuity with the knot of axons at the caudal end of the lesion makes it easy to identify regenerates for 0.5 cm caudal to the lesion (Fig. 4A). For a further 0.5 cm we saw a steadily decreasing number of labelled axons with the morphology of regenerates in the same position. We identified these as regenerated CST axons, although with lower confidence than for the more rostral 0.5 cm. In cross sections cca 1 cm below the lesion we observed a bundle of GFP labeled axons in intermediate and ventral gray matter in the immediate proximity to the central canal, which could represent some of the regenerating CST axons growing around the lesion ventrally (S. 2 A1, B1, Fig. 6).

**Fig. 4.**
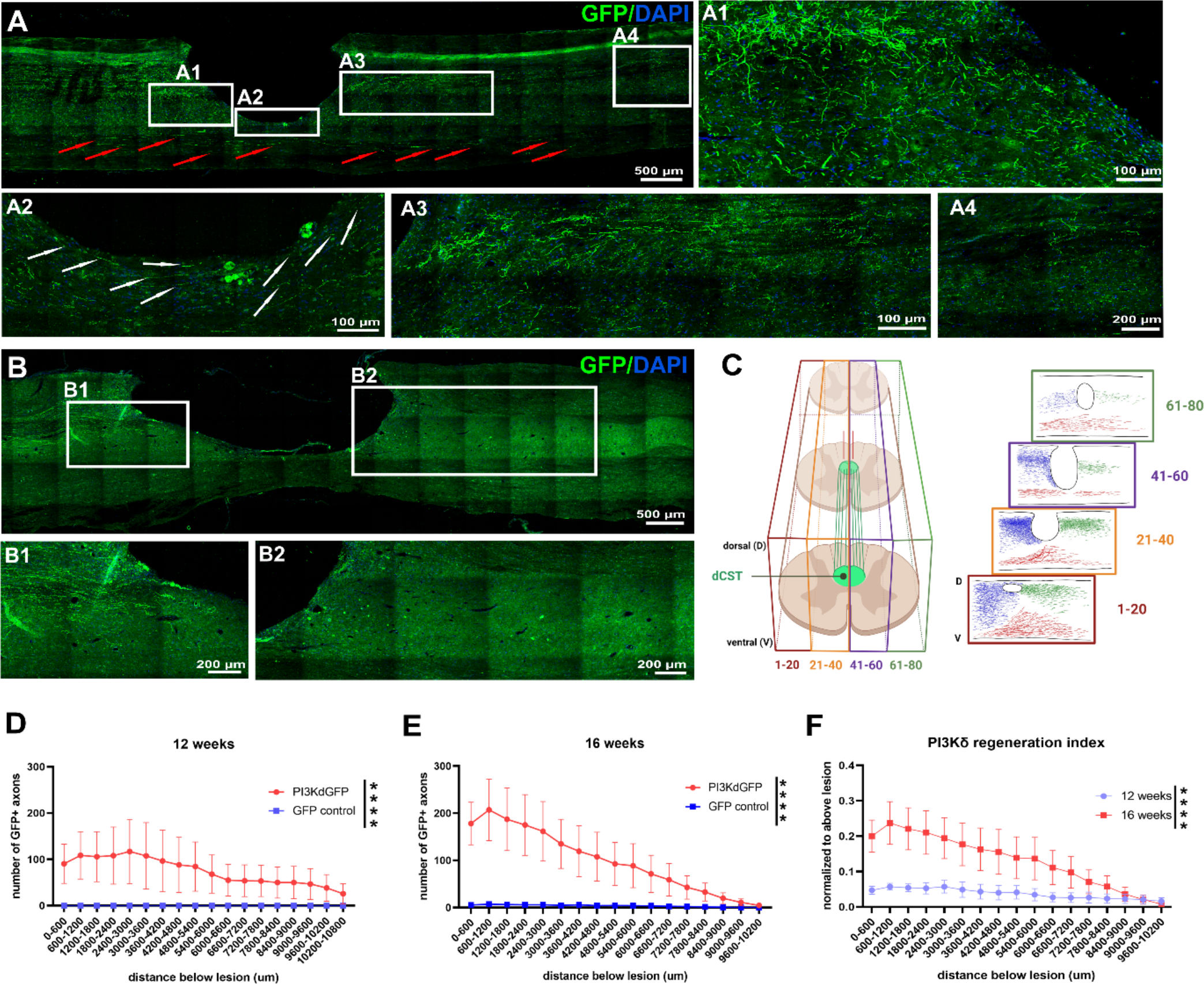
Axon count below lesion was evaluated in every other 20 μm spinal cord sagittal section in treated (A) and control animals (B). Sums of axons were calculated for each 600 μm and their distribution up to 1 cm depicted (D). Regeneration indices were determined representing axon count normalized to values above lesion and compared between 12 and 16 weeks after SCI and treatment (E). A reconstruction of axon growth in the lesion area and below was drawn using Biorender (C). Drawing was inspired by [20]. Data shown as mean and SEM, paired t test, **** *p<0*.*0001*

Additionally, we reconstructed the growth pattern in AAV1-CAG-PIK3CD treated rats that were sectioned frontally and confirmed, that axons also have the tendency to bypass the lesion laterally at 9 and 12 weeks after injury, and more are found when allowed additional time (Fig. 5C).

**Fig. 5.**
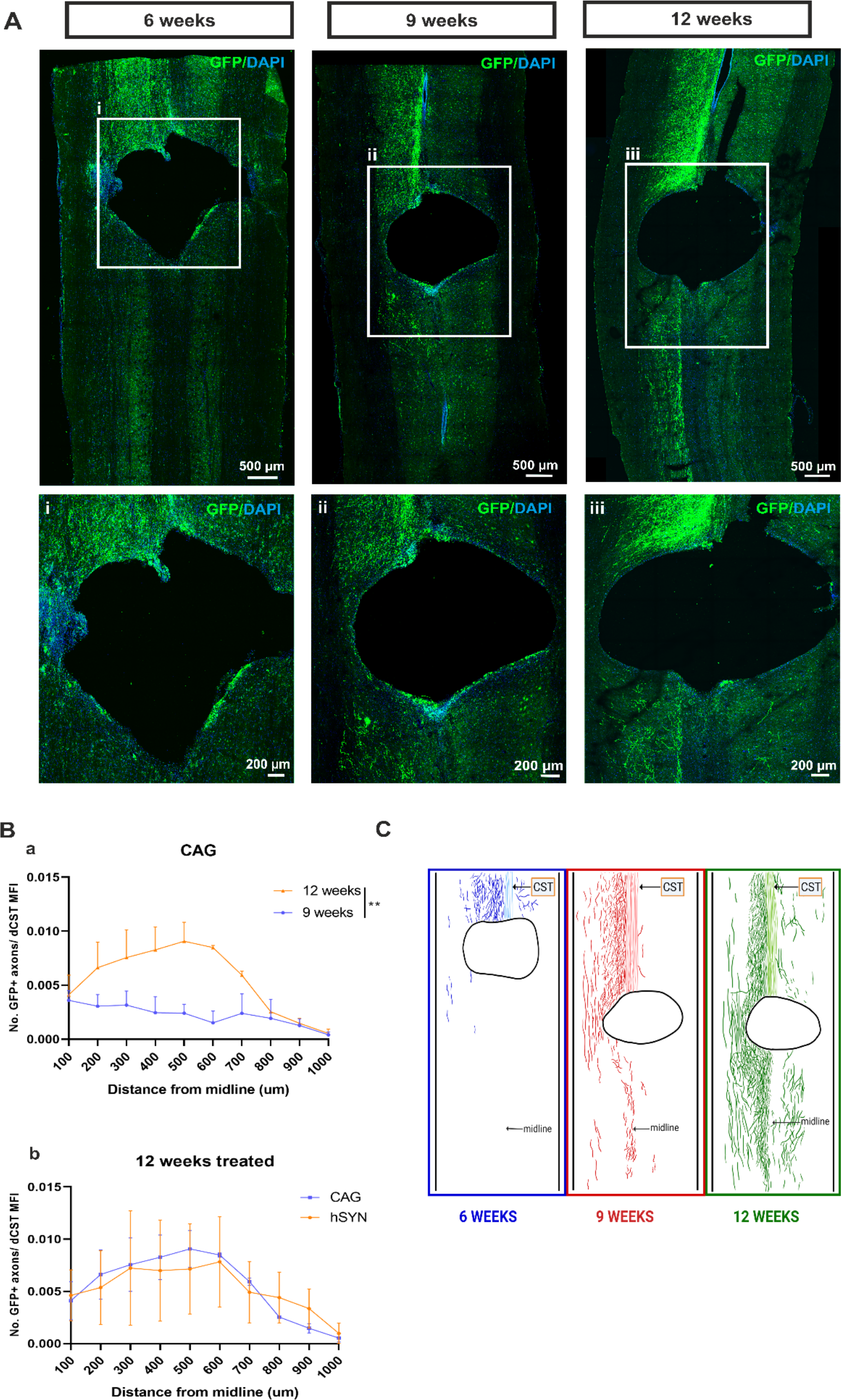
Frontal plane of sectioning spinal cords from rats treated with AAV1-CAG-PIK3CD + AAV1-CAG-eGFP vector mixture at 6, 9 and 12 weeks post injury and cortical injections. Axons reach the lesion border and start bypassing the lesion laterally to the cavity and beyond (Ai, Aii, Aiii). Reconstruction of axon growth pattern (C) Sprouting comparison between CAG 9 wks and 12 wks show significant increase in sprouting at 12 weeks (Ba) and no difference between CAG and hSYN promoters at 12 weeks (Bb).

### GFP labeled axons below the lesion colocalize with excitatory presynaptic marker

We looked for presynaptic varicosities in the GFP labeled axons around 1 cm below injury identifying punctae of vGlut1/2, a marker of excitatory synapses, and GFP punctae. Many co-localized puncta were seen in spinal cord gray matter. (Fig. 6A, B, C, V. 1 – video, S. 2). This suggests, that newly formed axons may be forming synapses with targets below the lesion and contribute to functional improvements.

**Fig. 6.**
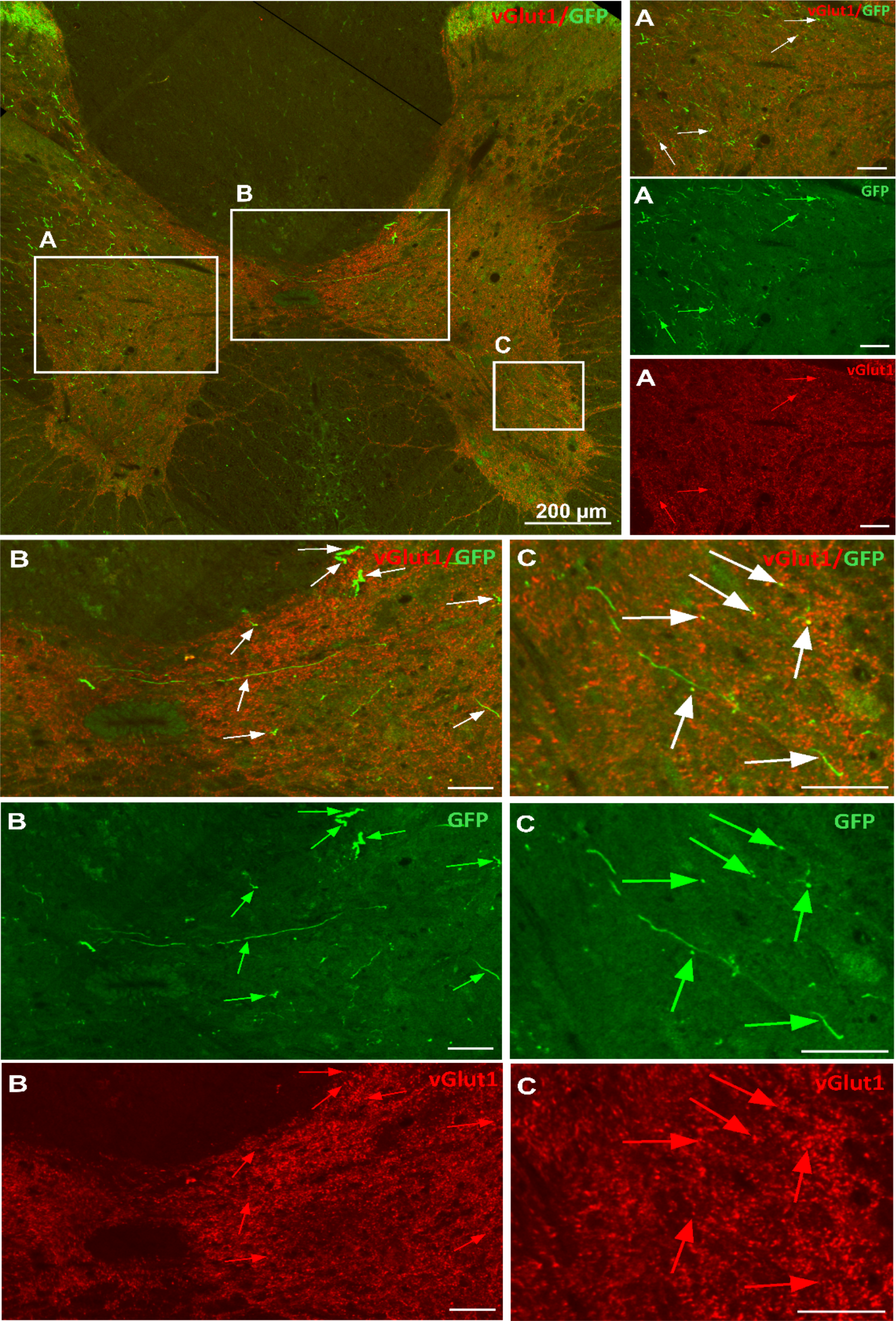
Synapse formation below lesion. In 20 μm cross sections, GFP+ axons below lesion colocalize with a vGlut1/2 excitatory synapse markers. Examples shown in ventral horn areas (A, C) and near the central canal (B). Images were acquired with Dragonfly spinning disc microscope at 40x magnification. Scale bars 50 μm unless otherwise stated.

### Overexpressing PI3Kδ improved behavioral outcomes and was motor function specific

Treated and control rats were behaviorally tested once per week for 16 weeks after injury following a 1-week recovery break. Motor tests included horizontal ladder crossing, grip strength and skilled paw reaching using the Montoya staircase. Motor recovery started at week 6 and gradually, treated rats outperformed controls in all motor tests. The best performances were recorded in the last period (of 6 – 4 weeks) of testing (Fig. 7A, B, C, D). Skilled paw reaching was significantly better in the left paw, however, we also recorded improvement in the right paw (Fig. 7C, D) in the number of pellets eaten as well as distance reached (data not shown). We included a sensory von Frey test to rule out hyperalgesia and saw very slight and insignificant improvement in sensation with time, which was still far below baseline. This confirmed that our treatment strategy was motor pathway specific and the minimal improvement was likely due to spontaneous recovery (Fig. 7E, F).

**Fig. 7.**
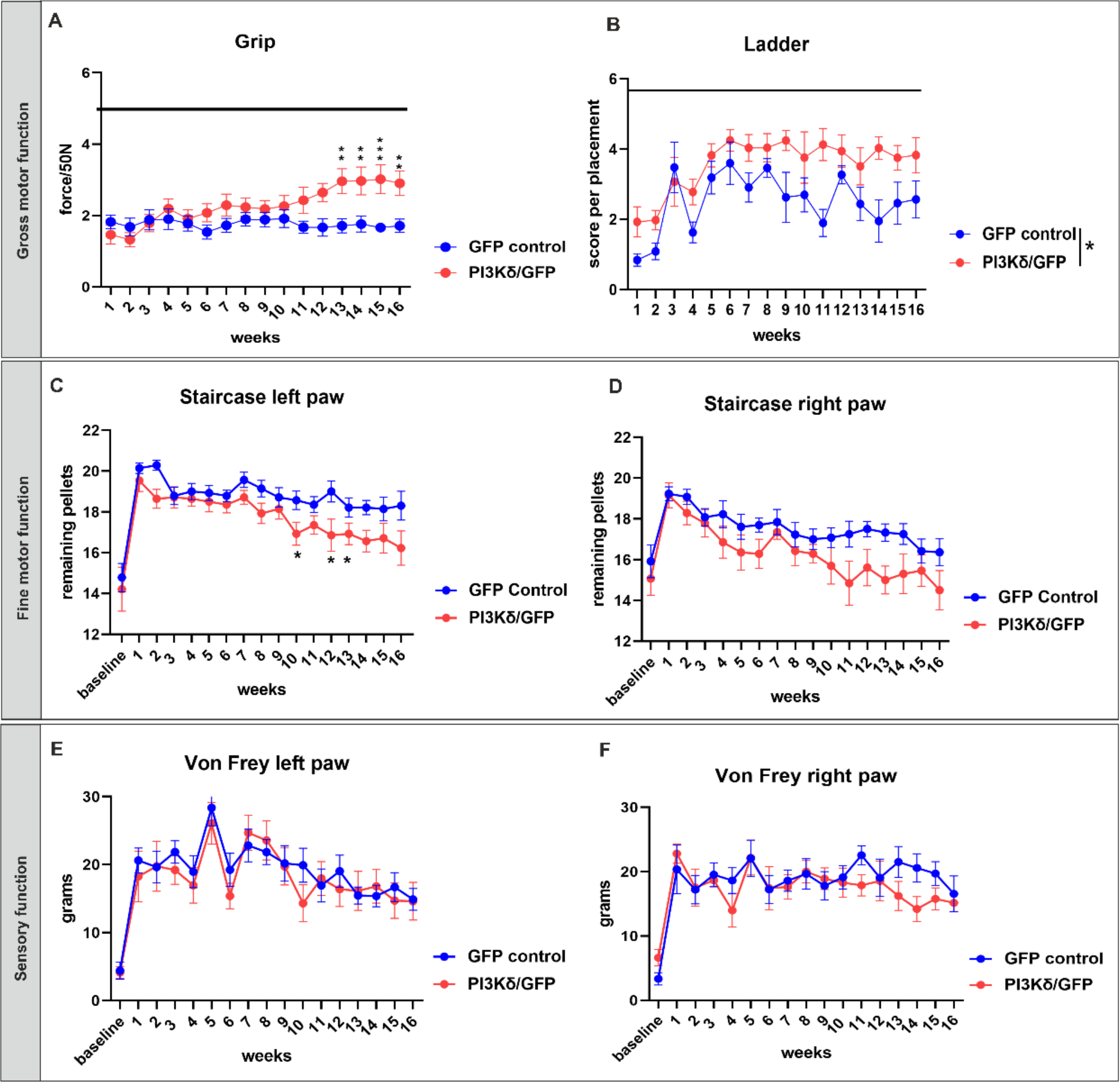
Lister Hooded rats were trained for 2 weeks and baseline was recorded, followed by lesion induction and cortical injections of vectors. After 1 week break, rats were tested once a week for 16 weeks to determine changes in gross (A, B) and fine (C, D) motor function. Rats expressing PI3Kδ (n=15) outperformed GFP controls (n=14) in grip strength, pellet reaching and ladder crossing. In grip assessment, Mixed model Two-way ANOVA revealed ** overall significance (*p = 0*.*002*) with Bonferroni post-tests significant differences at weeks 13 - 16 (A). Ladder crossings were scored on a scale 0 - 6, which captures errors in front paw placements. Mixed model Two-way ANOVA revealed * significant difference in performance between treated and control rats (*p=0*.*0295*) (B). Mixed model Two-way ANOVA and t-test were used to calculate * overall significance in skilled left paw reaching (*p = 0*.*0297*) with differences in individual weeks 10, 12 and 13 (C). Assessment of the right paw fine motor movement didn’t reach significant difference with *p = 0*.*0514* (D). Sensory von Frey test shows no difference between groups (E, F). Data shown as mean and SEM.

### Electrophysiology recordings confirmed functional improvements

In order to assess electrophysiological connectivity, spinal cord dorsal surface responses were measured in a different set of animals, as in previous studies [21], 1 mm laterally from the midline 1 cm above the lesion (Fig. 8B, S. 3A) and 1 cm below (Fig. 8A, B, C) in treated, uninjured and control animals 16 weeks after treatment. The right pyramid was stimulated by implantation of a tungsten needle electrode with increasing current amplitudes and cord dorsal potentials were measured with a silver ball electrode. After recordings, we confirmed that re-lesioning the cord lead to loss of signal. Significantly higher potentials were measured in AAV1-hSYN-PIK3CD treated rats when compared with their AAV1-hSYN-GFP treated controls but were not significantly different from uninjured animals (Fig. 8C, D). In addition to CDPs, we also measured EMG potentials in distal forepaw muscles of both left and right paws through transcutaneous needle electrodes. We recorded significantly greater responses in both contralateral and ipsilateral paws to the treated hemispheres, further corroborating our axon sprouting analysis results (Fig. 8E, Fig. 3).

**Fig. 8.**
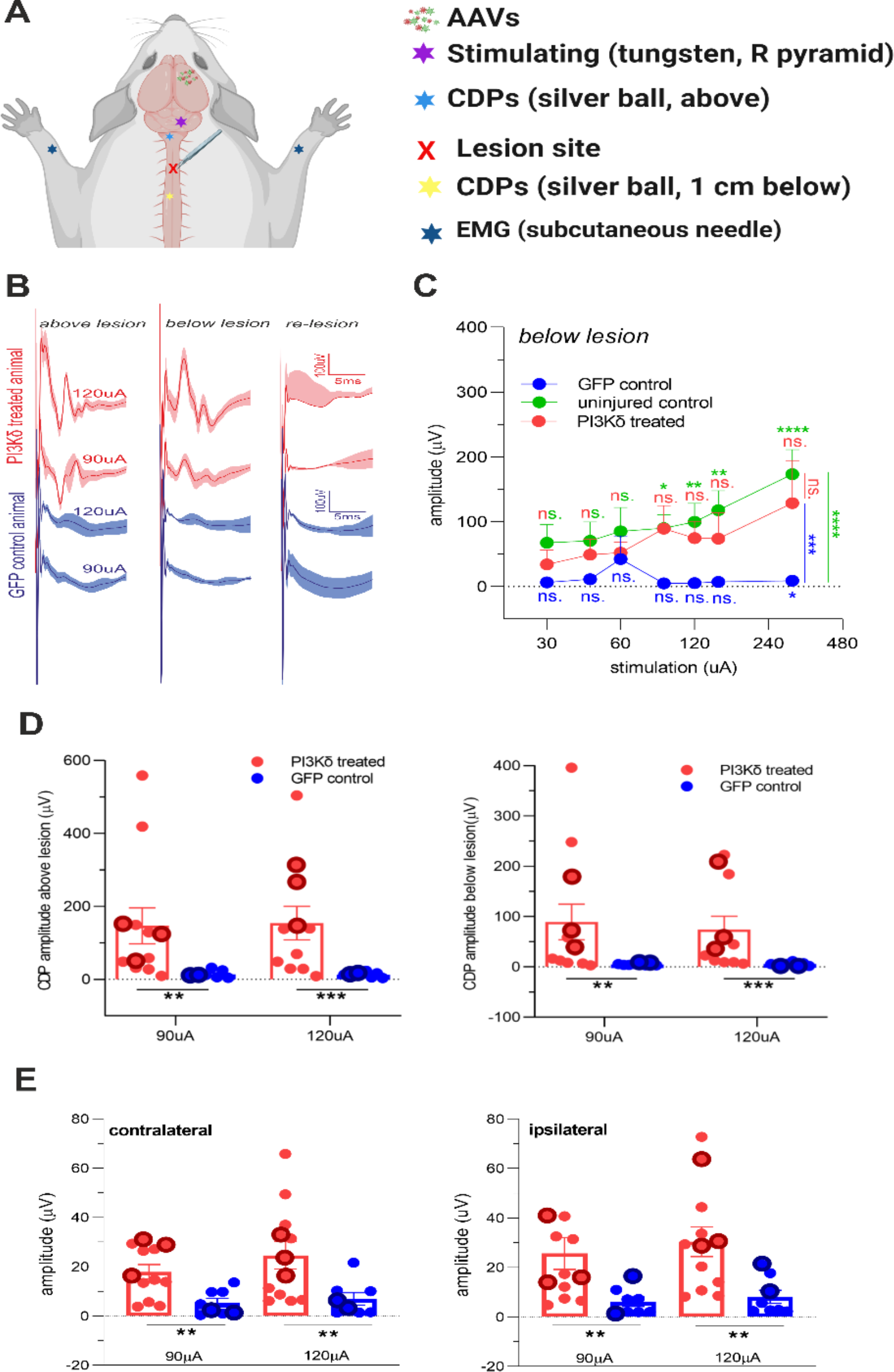
Functional recovery of PIK3CD treated rats (n=12) was confirmed with electrophysiological recordings, in which the right pyramid was stimulated using a tungsten electrode (5 square wave pulses at 300 Hz) with increasing current amplitudes (30 - 300 μA). CDPs were measured on spinal surface laterally to midline with silver ball electrodes above and below lesion. EMGs of distal forepaw muscles were recorded concurrently (A). Examples of CDP responses in treated and control rats at 90 μA and 120 μA (B). Recordings were compared with healthy uninjured rats (n=6) and GFP control rats (n=9). Significant differences were found between groups with two-way ANOVA and Sidak’s post-tests. No significant difference (ns) between uninjured and PIK3CD treated rats was found 1 cm below lesion as opposed to controls, which elicited significantly smaller responses than both treated and uninjured rats (C). Significantly stronger responses were recorded in PIK3CD treated rats compared to GFP controls at 90 μA and 120 μA stimulating currents (D). EMG recordings in distal flexor muscles show significantly better responses at 90 μA and 120 μA stimulating currents (E). Data shown as mean and SEM. *\*p<0*.*05; *** p<0*.*001; **** p<0*.*0001*

### Neuronal activity in spinal neurons was upregulated after stimulation of PI3Kδ treated rats

To see whether electric stimulation elicited a molecular response indicative of neuronal activity in postsynaptic spinal cord neurons, expression of the early immediate gene neuronal activity marker cFOS was evaluated. After recordings, animals were kept for additional 2.5 hours before perfusion so that cFOS analysis could be carried out in the areas above (S. 3) and below lesion (Fig. 9). Numbers of cFOS positive nuclei were counted around the central canal, in both ventral horns and in both dorsal horns (Fig. 9A). Increased expression density of nuclear cFOS was determined in both areas in treated rats. Significant differences were found in ventral and dorsal horns contralaterally to the treated hemisphere in AAV1-hSYN-PIK3CD animals when compared to their controls, but not when compared to uninjured rats (Fig. 9E, F).

**Fig. 9.**
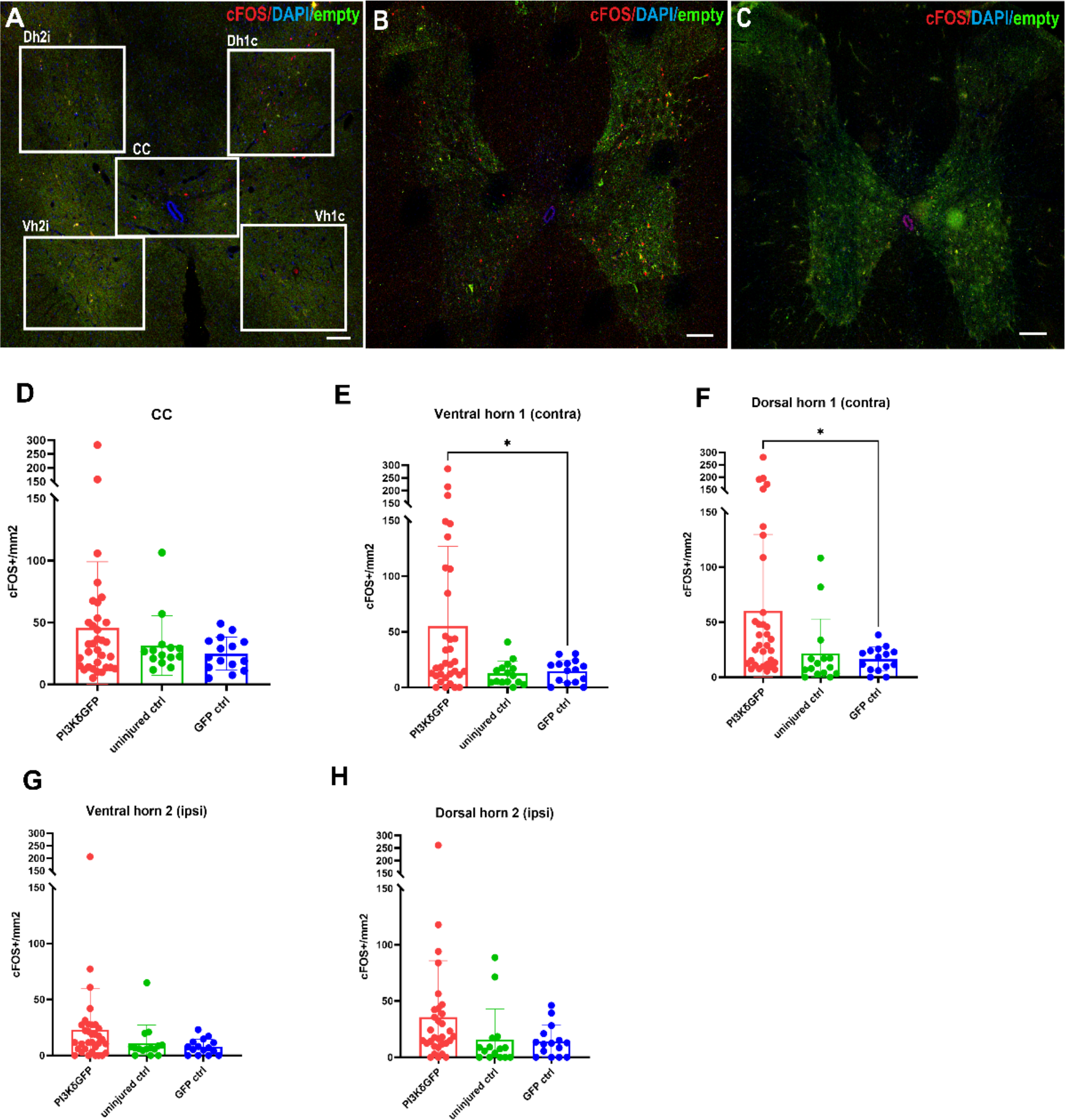
cFOS+ nuclei were counted in 5 regions of 20 μm spinal cross sections 1 cm below lesion. Representative images show cFOS activation pattern in stimulated uninjured (A), PIK3CD treated (B) and GFP control rats (C). No statistical difference was found in the area adjacent to central canal (CC) between treated (n=11), control rats (n=5) and uninjured shams (D). Differences were determined with One-way ANOVA in both ventral (Vh1c) and dorsal horns (Dh1c) (E, F) where the density of cFOS+ nuclei is higher in treated rats as opposed to GFP controls (*p=0*.*433* and *p=0*.*0288*, respectively). No statistically significant difference was found in the ipsilateral side (G, H). Data shown as mean and SEM. Scale bars 200 μm.

## Discussion

The aim of the study was to test the ability of a hyperactive form of PI3K, PI3Kδ, to enable regeneration of corticospinal tract (CST) axons in the adult rat spinal cord, and the ability of these regenerated axons to restore sensorimotor function and physiological connectivity. PI3Kδ under a CAG or synapsin promotor in an AAV1 vector was expressed in many output neurons in the sensorimotor cortex. Increased S6 phosphorylation in these neurons, determined in the hSYN promoter arm of this study, indicated that the enzyme was activating downstream signaling. CST axons were traced by co-transfection of eGFP. By 6 weeks after a C4 dorsal column lesion CST axons were seen regenerating around the margins of the lesions, with numbers progressively increasing at 9, 12 and 16 weeks. By 16 weeks many axons were seen that had regenerated past the lesions and on down the cord, with numbers steadily decreasing to 1cm, which is mostly where our sagittal tissue sectioning reached. Even beyond this distance, we saw GFP labeled axons in spinal cord cross sections, which associated with a presynaptic excitatory vGlut1/2. In these sections we found a bundle of axons right below the central canal in the gray matter, which could be some of the ventrally growing regenerates. Treated animals recovered in forepaw muscular strength, skilled paw reaching and ladder walking tasks, and electrical connectivity from the pyramids to below the lesions could be seen in cord dorsal electrode and distal forelimb EMG recordings. This electrical stimulation led to an increase in cFOS nuclear signal in the spinal cord gray matter both above and below the lesion site. Overall, the study shows that PI3Kδ expression stimulates extensive CST regeneration, electrophysiological connectivity and functional recovery.

### Axon regeneration in this study

Regenerating axons in the present study were seen mainly in regions surrounding the lesions. In general, they followed the edge of the lesions ventrally and laterally, usually within 200 μm. Having passed around the lesion cavities, axons collected at their caudal edge, where many fine axons with the typical fine meandering anatomy of regenerated axons could be found randomly oriented. From here, most of the axons grew in a more directed fashion in a caudal direction where a band of axons were seen within approx. 0.25 mm from the dorsal surface, their number steadily diminishing with distance from the lesion. Our lesions were dorsal, so the ventral CST was not cut. Thick straight unlesioned axons were therefore seen in the ventral cord when sectioned sagittally. Between this group of clearly unlesioned axons and the clearly regenerated axons there is a region of grey matter containing many fine processes, which could be branches from the regenerated axons or the unlesioned ventral CST; neurites in this region were not counted when they were seen to emerge from the ventral CST. The groups of processes in one of the rats are demonstrated in different colours in Figure 4.

### PI3K signaling and axons

PI3K and PIP3 signaling has been associated with axon growth in many studies, while inhibitors of PI3K are blockers of axon growth. Raised PIP3 levels have been implicated in CNS axon regeneration in many studies in which PTEN has been deleted or inhibited [22]. The action of PTEN is to dephosphorylate PIP3 back to PIP2, so reversing the action of PI3K. PTEN knockout therefore prevents PIP3 dephosphorylation, thus increasing its level. PTEN deletion has been shown to increase regeneration of axons in the optic nerve, CST and other CNS pathways. However, the regeneration-inducing effect of PTEN deletion falls off with age [23]. *In vitro* studies with PIP3 staining have shown that PIP3 levels in the processes of cortical neurons fall with age, and PTEN knockdown has little effect at increasing them [4]. Neurons contain plentiful PI3Kα [4], implying that there is little activation of this enzyme in mature neurons and therefore little PIP3 generation. PTEN deletion in the absence of PIP3 generation will therefore have little effect. For this reason, a hyperactive form of PI3K, PI3Kδ, was transduced into cortical neurons. A previous study has shown that this form of the enzyme raises PIP3 levels and the probability of axon regeneration in mature cortical neurons *in vitro* [4]. PIP3 is generally considered to signal through the AKT pathway, with many downstream targets, in particular protein translation via mTOR. It is probable that local translation of mRNAs in axons is increased by PIP3/AKT signaling, with positive effects on axon regeneration. However, PIP3 has widespread effects on proteins with PH and FYVE domains. These include several GAPs and GEFs that control axonal transport and trafficking [3]. Upregulation of neuronal mTORC1 complex, activated specifically by Akt1 or 2, leads to an increase in palladin levels concurrently with upregulated S6 phosphorylation. This cytoskeleton modulating protein regulates developmental axon extension and localizes in cortical neuron cell body, but at lower degree when compared to axon and growth cone domains. Increased translation of palladin results in the formation of multiple axons in mTORC1 dependent fashion. Additionally, authors of this study presented evidence of mTOR dependent local palladin translation within studied neurites [24]. Not all members of the Akt/mTOR pathway seem to have beneficial role in regeneration after SCI when upregulated. A study where neuronal Akt3 was strongly overexpressed with high titre gene therapy lead to seizures that were associated with increased size of neuronal cell body, which lead to megalencephaly of the injected hemisphere. This was prevented when authors adjusted the AAV titre. Robust new growth was seen, albeit without functional motor recovery [25]. Overexpression of PI3Kδ in this study did not lead to seizures or aberrant macro – or microscopic morphology.

### *PI3K*δ *and CST regeneration*

PI3Kδ is a hyperactive form of PI3K. It behaves very similarly to PI3Kα that has a mutation H1047R. These mutations mimic and enhance dynamic events in the natural activation process of the autoinhibited p85–p110 heterodimer, reviewed in[26], with the result that the enzyme is more readily activated and generates more PIP3 [27]. Expression of PI3Kδ in mature cortical neurons restores PIP3 levels in growth cones and enables their axons to regenerate after laser axotomy. Expression in retinal ganglion cells stimulates axon regeneration after optic nerve crush in both transgenic mice and AAV2 mediated delivery approach [4].

How might PI3Kδ expression cause the CST regeneration that we observed? PIP3 signals through several pathways, so its effects on axon regeneration are various. Local translation of axonal mRNAs is important for efficient regeneration. It is probable that signaling via AKT and mTOR will promote this process. Some key cell surface receptors such as integrins travel into axons in Rab11 recycling endosomes, and these are found in the cell body and dendrites but are excluded from axons by changes in axonal transport [28,29], with Rab11/integrin transport becoming retrograde to remove the items from axons. *In vitro* experiments showed that expression of hyperactive PI3Kδ restores anterograde transport of Rab11 vesicles and integrins into mature axons and promotes strong axon regeneration of cortical neurons [4].

In a previous study, we showed that PI3Kδ had effects on axonal transport, promoting transport of Rab11 recycling endosomes and integrins [3], probably through effects on RAB GAPs and GEFs. Several other receptors are also transported in Rab11 endosomes, so PI3Kδ treatment will have allowed transport of several molecules involved in regeneration. In addition, through actions on adaptors and scaffolds there will be effects on cytoskeletal dynamics.

### Increased plasticity, sprouting and branching and extrapyramidal pathways

In this study, we show that AAV assisted overexpression of PI3K delivered into rat motor cortex resulted in functional recovery after SCI. Treated rats showed robust GFP labeling of CST axons above the C4 lesion, in its vicinity ventrally and laterally as well as below the lesion reaching impressive distance. We acknowledge that with our model it is challenging to determine the proportions between true regeneration, sprouting over both short and long distances, and generation of new compensatory collaterals from the intact ventral CST. We believe that all of these types of axonal growth were part of the PI3Kδ effect.

Looking at cross sections above the C4 lesion, we were able to see clear labeling of the dorsal as well as ventral and lateral CST. Our axon sprouting analysis (Fig. 3) showed much increased axon sprouting of the dCST with many axons crossing the midline and reaching predominantly the ventral horns ipsilateral to the injected hemisphere. Moreover, we took notice of some sprouts from the ventral CST, which comprises a small but functionally competent part of this descending tract [30]. Growth of axons in studies enabling CST regeneration is generally seen mainly if not solely in the gray matter and our study is not an exception. It is therefore not impossible for these sprouted axons located in the gray matter and reaching ventral horns to continue growing through this area caudally to lesion site and beyond, starting to bypass it in this manner well above the lesion. The same could be true for the expectedly smaller portion of vCST sprouts. Such growth ipsilaterally to injected hemisphere could help explain the recorded EMGs in the ipsilateral paw (Fig. 8). Additionally, we cannot exclude the possible involvement of extrapyramidal tracts, such as the rubrospinal or reticulospinal. The reticulospinal tract descends ipsilaterally and innervates the trunk muscles and helps with weight support. We did notice, without further analysis, that treated rats supported their weight more readily and for prolonged time, which would be in agreement with this speculation. Moreover, this tract along with the rubrospinal tract could also be a source of compensatory collaterals.

In trying to understand the newly formed CST tract and its reorganization, we provide evidence that PI3Kδ overexpression does stimulate the regrowth of severed axons as we have seen them in sagittal and frontal sections progressively growing around the lesion both ventrally and laterally (Fig. 4, Fig. 5). This can be seen in cross sections from cca 1 cm below the C4 lesion. We can notice a bundle of axons right below or at the central canal caudoventrally to the lesion. It is possible that these are some of the truly regenerated axons (S. 2).

We observed noticeable branching of axons making it more complicated to distinguish whether they are branched sprouts, collaterals from spared parts of CST or branched true regenerates. This is a significant weakness of this study, and we acknowledge that better imaging techniques like light sheet microscopy would provide us with this information.

### Perspectives

PI3Kδ treatment enabled CST regeneration, but the length of robust regeneration was shorter than the 4-6 cm of a typical human spinal cord injury; this extent of regeneration is a target for translation to human therapy. In order to further increase regeneration PI3Kδ treatment could be combined with one or more stimulators of regeneration. A successful regeneration promotor for sensory axons in the spinal cord, able to achieve regeneration of over 4 cm, is integrin α9 combined with its activator kindlin-1. In sensory axons, which transport integrins, the combination of this tenascin-C-binding integrin and the kindlin activator led to extensive regeneration for several centimeters (Stepankova et al., 2023). Because PI3Kδ restores integrin anterograde transport in cortical neurons, combining it with integrin could be successful [4,28]. PI3Kδ expression led to upregulation of S6 phosphorylation which mediates feedback inhibition of the AKT/mTOR pathway. There is, however, evidence that an inhibitor of S6 phosphorylation, via S6K1, is a promoter of CST regeneration [31]. This group confirmed that regenerative effects were mediated by the PI3K/mTOR pathway with negative feedback provided by S6K1. A combination of PI3Kδ with an inhibitor of S6 phosphorylation is therefore a potentially successful combination.

## Supplementary material legends

**S**.**1** Lesion completeness was confirmed with either PKCγ staining (Wistar) of the dorsal CST above (A) and below (B) lesion, or by an MRI scan of the whole cord (Lister Hooded) in both PI3Kδ treated (C) or control rats (D). Scale bars 100 μm.

**S. 2** Cross sections from 2 different rats (A, B) showing the central canal area (A1, B1) both with axons in the gray matter right below the central canal approximately 1 cm caudally to the lesion border. These axons are annotated with white arrows.

**S. 3** CDP measurements from above the lesion were recorded and used as guidance of proper electrode placement. All measured rats elicited responses from this area, but they were significantly smaller in controls (A). cFOS^+^ nuclei densities in spinal cord gray matter shows higher values of neural activation in AAV1-hSYN-PIK3CD treated rats around central canal (B), contralateral ventral and dorsal horns (C) and in ipsilateral dorsal horn (D) when compared to AAV1-hSYN-eGFP controls.

**V .1** Imaris rendered 3D reconstruction of Fig. 6B where colocalizations of GFP and vGlut1/2 without potential 2D artifacts can be better appreciated.

## Acknowledgement

This research was supported by the International Foundation for Research in Paraplegia **(IRP) P186, OPJAK CZ**.**02**.**01**.**01/00/22_008/0004562** and by the Microscopy Service Centre of the Institute of Experimental Medicine CAS supported by the **MEYS CR** (**LM2023050 Czech-Bioimaging**), which includes support by the Ministry of Education, Youth and Sports of the Czech Republic (**Research Infrastructure NanoEnviCZ, LM2018124**) and by The European Union – European Structural and Investment Funds in the frame of the Research Development and Education – project Pro-NanoEnviCZ operational program (**Project No. CZ**.**02**.**1**.**01/0**.**0/0**.**0/16_013/0001821**).

