## Supplementary figures and images for "Hyperactive PI3Kinase delta enables long distance regeneration of the rat corticospinal tract"

### S. 1

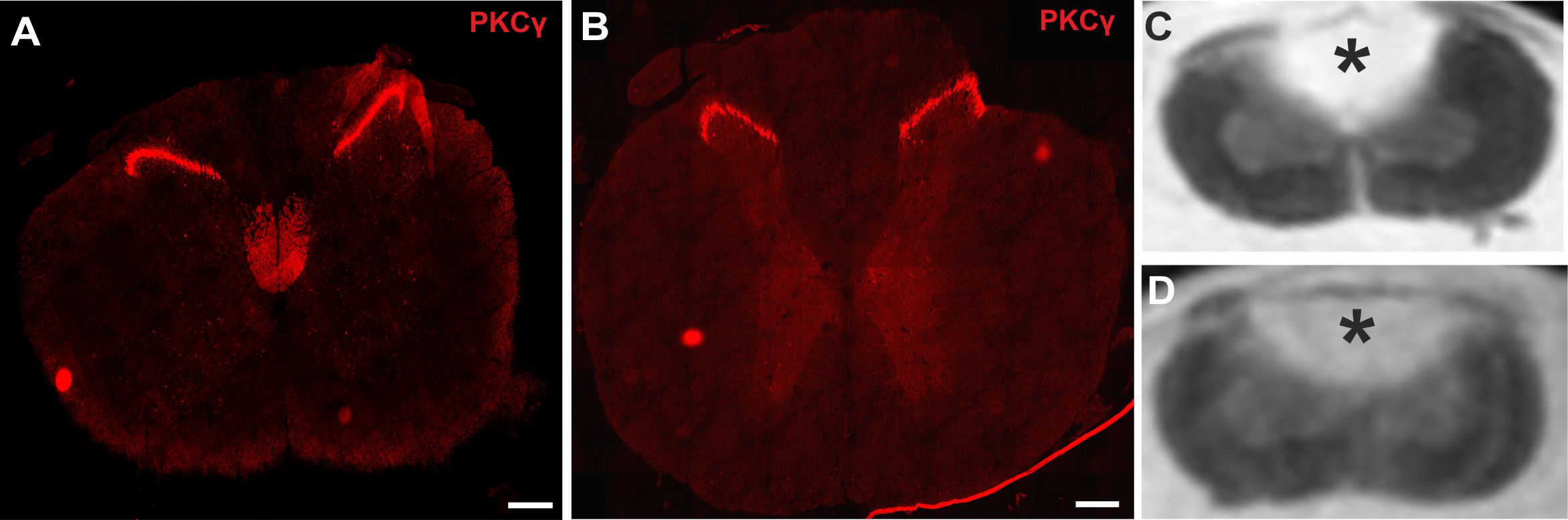

### S. 2

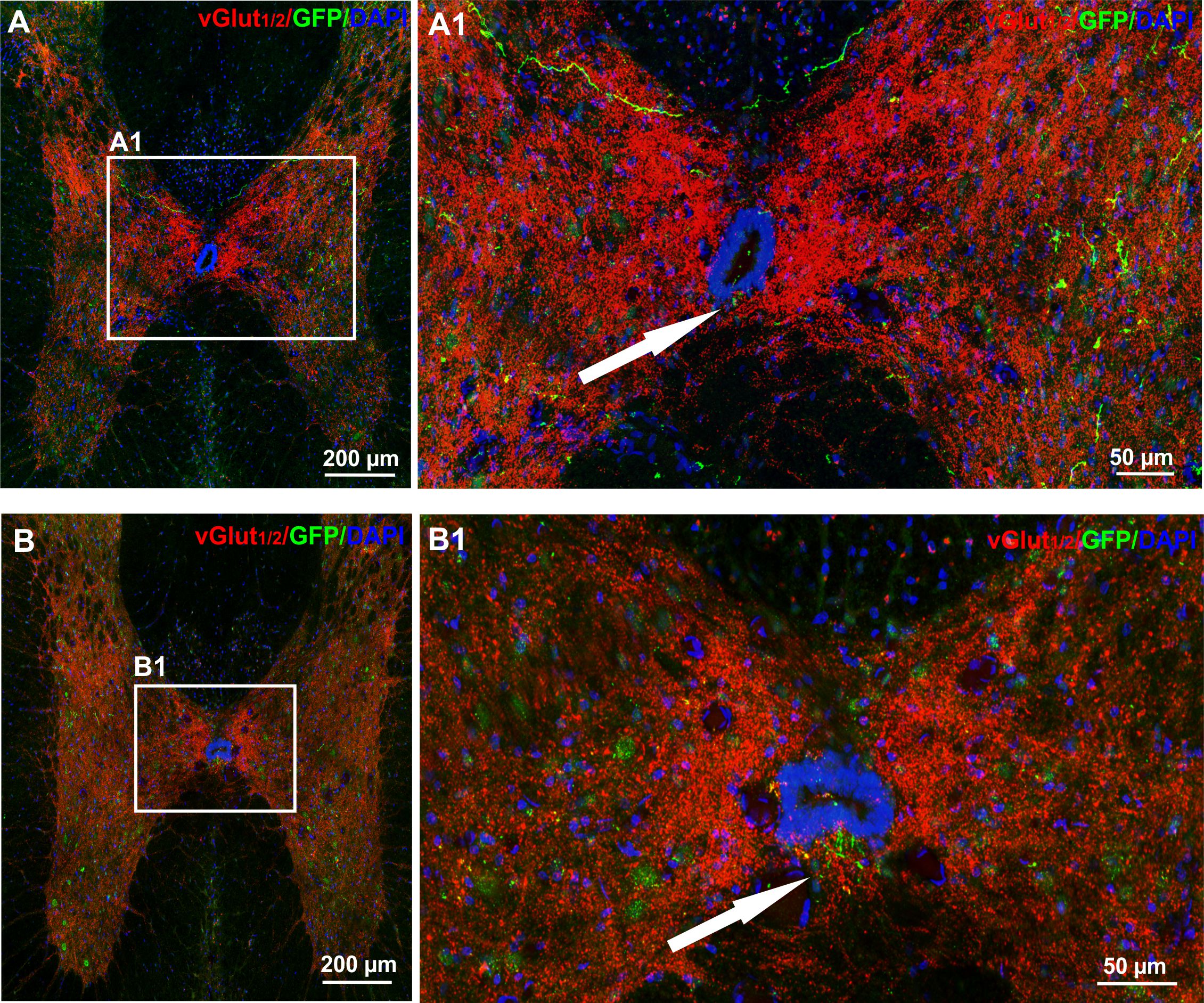

### S. 3

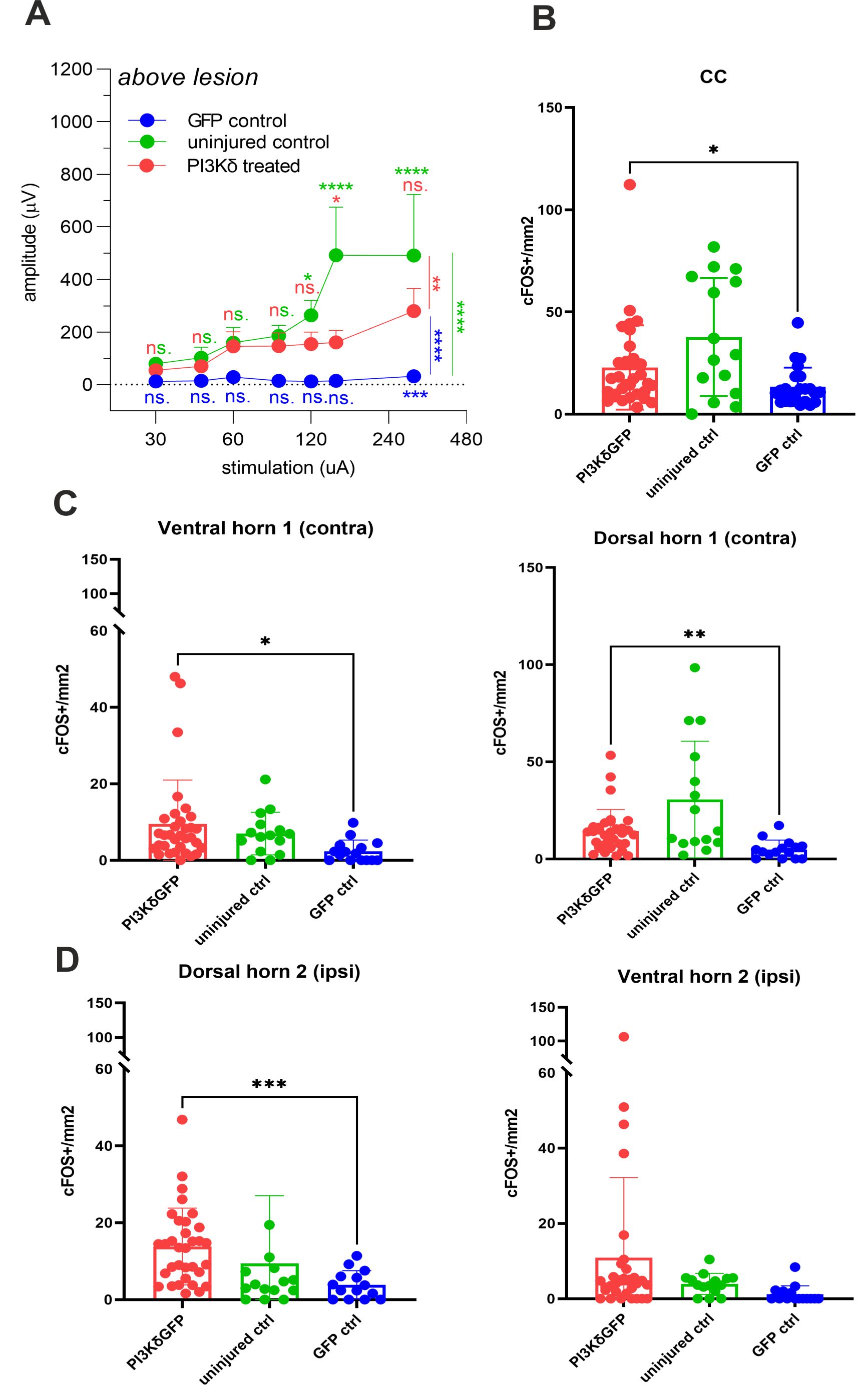
